# A chiral selectivity relaxed paralog of DTD for proofreading tRNA mischarging in Animalia

**DOI:** 10.1101/123919

**Authors:** Santosh Kumar Kuncha, Mohd Mazeed,, Raghvendra Singh,, Bhavita Kattula,, Satya Brata Routh,, Rajan Sankaranarayanan

## Abstract

D-aminoacyl-tRNA deacylase (DTD), a *trans*-editing factor found in bacteria and eukaryotes, removes D-amino acids mischarged on tRNAs as well as achiral glycine mischarged on tRNA^Ala^. An invariant cross-subunit Gly-*cis*Pro motif forms the mechanistic basis of strict L-amino acid rejection from the catalytic site. Here, we present the identification of a DTD variant, named ATD (<u>A</u>nimalia-specific <u>t</u>RNA <u>d</u>eacylase), that harbors a Gly-*trans*Pro motif. The *cis*-to-*trans* switch causes a “gain of function” through L-chiral selectivity in ATD resulting in the clearing of L-alanine mischarged on tRNA^Thr^(G4•U69) by eukaryotic AlaRS. The biochemical proofreading activity of ATD is conserved across diverse classes of phylum Chordata. Animalia genomes enriched in tRNA^Thr^(G4•U69) genes are in strict association with the presence of ATD, underlining the mandatory requirement of a dedicated factor to proofread tRNA misaminoacylation. The study highlights the emergence of ATD during genome expansion as a key event associated with the evolution of Animalia.

## INTRODUCTION

Translational quality control is a complex and tightly regulated process which involves editing of errors in most scenarios. However, it also encompasses a targeted and selective compromise in fidelity, thereby allowing percolation of errors under specific conditions such as oxidative stress. It ensures an optimum dynamic balance in the cellular proteome and hence overall cellular homeostasis. A multitude of factors—from aminoacyl-tRNA synthetases (aaRSs) to ribosome as well as proteasome—play significant roles in performing this complex phenomenon (**Brandman and Hegde, 2016; Bullwinkle et al., 2014; Guo and Schimmel, 2012; Ibba and Söll, 2000; Moghul et al., 2014; Ogle and Ramakrishnan, 2005; Pouplana et al., 2014; Rodnina, 2016; Rodnina and Wintermeyer, 2016; Schwartz and Pan, 2017; Simms et al., 2017**). A key step in this process includes decoupling of D-amino acids mischarged on tRNAs. This function, termed “chiral proofreading”, is performed by a dedicated *trans*-editing factor called Daminoacyl-tRNA deacylase (DTD) (**Ahmad et al., 2013; Calendar and Berg, 1967; Ferri-Fioni et al., 2001; Soutourina et al., 1999**). The chiral proofreading enzyme forms one of the major cellular checkpoints, which also includes aaRSs, elongation factor Tu (EF-Tu) and ribosome, to prevent infiltration of D-amino acids into translational machinery (**Agmon et al., 2004; Ban et al., 2000; Bhuta et al., 1981; Englander et al., 2015; Jonak et al., 1980; Ling et al., 2009; Pingoud and Urbanke, 1980; Yamane et al., 1981**).

DTD—present throughout Bacteria (except cyanobacteria) and Eukarya—employs an invariant cross-subunit Gly-*cis*Pro motif in the active site to ensure substrate stereospecificity. The *cis* conformation of the motif disposes the two carbonyl oxygens in a parallel orientation, projecting them directly into the active site pocket (**Ahmad et al., 2013**). Such an architecture of DTD’s chiral proofreading site leads to steric exclusion of even the smallest amino acid with L-chirality, *viz.*, L-alanine. Thus, strict L-chiral rejection forms the only mechanistic basis of DTD’s enantioselectivity. Consequently, the chiral proofreading site is completely porous to achiral glycine and exhibits unwarranted activity on the cognate Gly-tRNA^Gly^. The glycine “misediting paradox” thereby generated is effectively resolved through protection of the cognate achiral substrate by EF-Tu (**Routh et al., 2016**). Nevertheless, our recent findings have demonstrated that the porosity of DTD’s active site to glycine is advantageous, since it enables the enzyme to efficiently clear the non-cognate Gly-tRNA^Ala^ species generated by alanyl-tRNA synthetase (AlaRS). Therefore, DTD’s cellular function extends beyond just chiral proofreading during faithful translation of the genetic code (**Pawar et al., 2017**). Interestingly, in archaea and cyanobacteria, chiral proofreading is performed by DTD2 and DTD3, respectively; the latter two are non-homologous to DTD (**Ferri-Fioni et al., 2006; Wydau et al., 2009**). However, in archaea, which lack DTD, a DTD-like module is covalently appended to threonyl-tRNA synthetase (ThrRS) as the N-terminal domain (NTD) that edits L-serine misacylated on tRNA^Thr^ (**Dwivedi et al., 2005; Hussain et al., 2006, 2010; Korencic et al., 2004**). Thus, both DTD-like fold (comprising DTD and NTD) and chiral proofreading function (performed by DTD, DTD2 and DTD3) are conserved across all domains of life. Biochemically, the DTD-like fold is an RNA-based catalyst that employs only the 2′-OH of adenosine-76 (A76) at the 3′-terminus of tRNA rather than protein side chains for catalysis at the RNA–protein interface (**Ahmad et al., 2015; Routh et al., 2016**).

Proofreading during aminoacyl-tRNA synthesis has been proposed and extensively studied so far in the context of errors only in amino acid selection by aaRSs (**Dock-Bregeon et al., 2000; Fersht, 1977, 1998; Fukai et al., 2000; Jakubowski and Fersht, 1981; Lincecum et al., 2003; Matinis and Boniecki, 2010; Nureki et al., 1998; Pauling, 1958; Perona and Gruic-Sovulj, 2014; Silvian et al., 1999; Yadavalli and Ibba, 2012**). Defects in proofreading have been associated with multiple cellular pathologies including neurodegeneration in mouse and cell death (**Bacher et al., 2005; Bullwinkle et al., 2014; Bullwinkle and Ibba, 2016; Karkhanis et al., 2007; Kermgard et al., 2017; Korencic et al., 2004; Lee et al., 2006; Liu et al., 2014; Lu et al., 2014; Moghal et al., 2016; Mohler et al., 2017; Nangle et al., 2002; Roy et al., 2004**). Amino acids are substantially smaller in size compared to tRNAs, and are also similar in structure/chemistry in several cases. Consequently, errors in amino acid selection by synthetases are significantly higher (about one in 10^3^–10^2^) than the overall error observed during translation of the genetic code (about one in 10^4^–10^3^) (**Loftfield and Vanderjagt, 1972; Perona and Gruic-Sovulj, 2014; Yadavalli and Ibba, 2012**). Errors in tRNA selection, which either happen naturally and constitutively or are induced by environmental conditions (such as oxidative/temperature/antibiotic stress), have been noted in several instances, although such mistakes are not as common as those in amino acid selection (**Gomes et al., 2007; Netzer et al., 2009; Schwartz and Pan, 2016; Schwartz et al., 2016; Sheppard et al., 2008**). However, dedicated proofreading factors for correcting mistakes in tRNA selection have not been reported till date.

Here, we describe the identification and characterization of a novel DTD-like factor, named ATD (Animalia-specific tRNA deacylase). Present in kingdom Animalia and more specifically all across phylum Chordata, ATD proofreads a critical tRNA selection error made by AlaRS. An unprecedented switch from the chiral-selective “Gly-*cis*Pro” dipeptide in DTD to “Gly-*tgans*Pro” in ATD is the key to ATD’s gaining of L-chiral selectivity. This “gain of function” through relaxation of substrate chiral specificity underlies ATD’s unique capability of correcting the error in tRNA selection. The strict coexistence of the proofreading factor with the error-inducing tRNA species underlines its requirement for translational quality control in Animalia. Our study represents the first instance of a proofreading factor being identified as one that is responsible for the correction of an error in tRNA selection during translation of the genetic code.

## RESULTS

### Two characteristic sequence motifs in ATD show subtle differences from those in DTD

While performing protein BLAST–based *in silico* search for DTD sequences, we came across many sequences in the database which are annotated as probable DTD2. However, these sequences bear no sequence similarity to the canonical DTD2 present in Archaea. Therefore, we renamed this protein ATD (as explained later) to distinguish it from the canonical DTD2 and avoid confusion over nomenclature. Moreover, DTD and ATD share less than 30% sequence identity between them which is significantly lower than that between DTDs (>50%) or between ATDs (>45%) (**Figure 1—figure supplement 1**). Besides, ATD also does not show homology with DTD3. Multiple sequence alignment of ATD and DTD sequences showed that ATD has – PQATL– and –TNGPYTH– as signature motifs, which are similar to though distinct from the corresponding active site motifs in DTD, *viz.*, –SQFTL– and –NXGPVT–, respectively (**Figure 1A**). Strikingly, some of the key conserved residues near DTD’s active site, involved in a network of interactions and responsible for holding the Gly-*cis*Pro motif, are also different in ATD. The most notable among these is a highly conserved arginine in DTD (Arg7 in DTD from *Escherichia coli* (EcDTD) or *Plasmodium falciparum* (PfDTD)) which is replaced by a conserved glutamine in ATD (Gln16 in ATD from *Mus musculus* (MmATD)) (**Figure 1A**). Thus, comparative analysis of ATD and DTD sequences showed subtle variations in some of the key conserved residues present in and near the active site.

**Figure 1.**
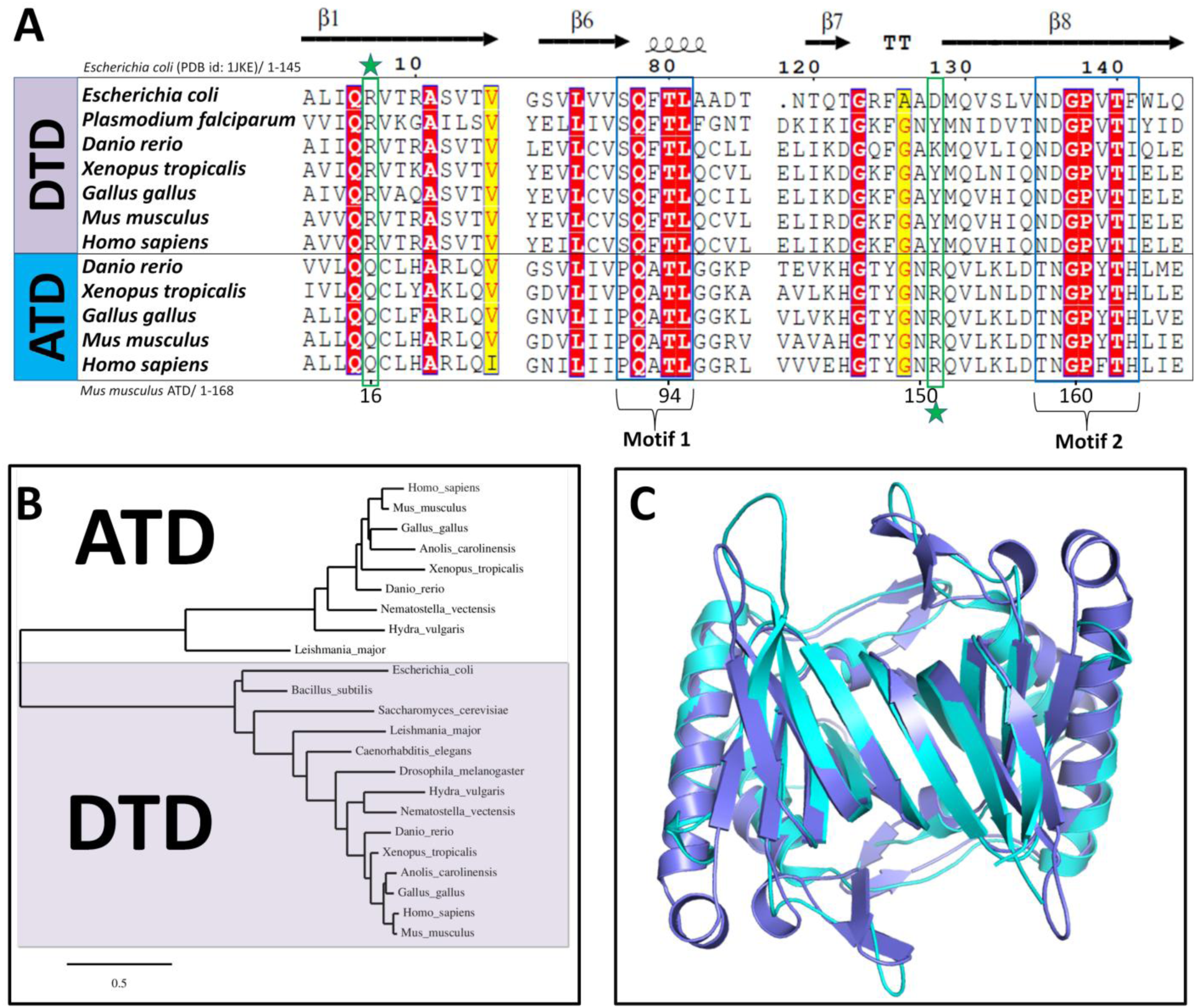
ATD is a variant of DTD. (**A**) Multiple sequence alignment showing similar but distinct and characteristic sequence motifs in DTD and ATD (motifs 1 and 2). The highly conserved arginine in DTD (Arg7, EcDTD) is indicated by a star above, whereas the invariant arginine in ATD (Arg151, MmATD) is highlighted by a star below. (**B**) Phylogenetic classification of DTD and ATD showing their grouping into two separate categories. (**C**) Crystal structure of MmATD homodimer (blue) superimposed on that of PfDTD homodimer (cyan; PDB id: 4NBI).

### ATD is found in kingdom Animalia and throughout phylum Chordata

A thorough *in silico* search for ATD sequences revealed that ATD is present in Eukarya, but absent in Bacteria and Archaea. Within Eukarya, ATD is present exclusively in kingdom Animalia, except for a few protozoa (four species of *Leishmania*, two of *Trypanosoma*, and one each of *Saprolegnia*, *Salpingoeca* and *Acanthamoeba*, whose genomes have been sequenced) (**Data 1**). More importantly, ATD is found all across phylum Chordata, whereas its distribution in non-chordate phyla is rather sparse (**Figure 1—figure supplement 2, Data 1**). It is worth noting that most of the protozoa that harbor ATD are parasites of various vertebrate hosts. Therefore, ATDs from these protozoa may be outliers as the possibility of horizontal transfer of ATD gene to these protozoa from their host organisms cannot be ruled out. Contrary to ATD’s restricted distribution in Animalia, DTD is found throughout Bacteria and Eukarya. Nevertheless, phylogenetic analysis of ATD and DTD showed that the two fall into two distinct groups (**Figure 1B**).

### ATD belongs to the DTD-like fold

To gain insights into ATD’s function, we solved the crystal structure of MmATD at 1.86 Å resolution. We were able to solve the structure by molecular replacement using PfDTD as the search model, despite the fact that the two share less than 30% sequence identity (**Figure 1C**, **Table S1**). Structural superposition of MmATD on PfDTD and NTD from *Pyrococcus abyssi* (PabNTD) showed an r.m.s.d. of 1.68 Å over 141 Cα atoms and 3.34 Å over 77 Cα atoms, respectively (**Figure 1C**, **Figure 1—figure supplement 3A**). As is the case with DTD and NTD, ATD too is a homodimeric protein. Interestingly, a Dali-based PDB search for structural homologs of ATD identified a protein (ATD) from *Leishmania major* (LmATD) which is annotated as a probable eukaryotic DTD, and deposited by Structural Genomics of Pathogenic Protozoa Consortium (**Fan et al., 2008**). Structural superimposition of LmATD on MmATD gives an r.m.s.d. of 1.29 Å over 148 Cα atoms (**Figure 1—figure supplement 3B**). Thus, like DTD and NTD, ATD also belongs to the DTD-like fold.

### ATD harbors a Gly-***trans*Pro motif in the active site unlike DTD’s Gly-*cis*Pro motif**

The crystal structure of ATD revealed that its characteristic motifs (–PQATL– and –TNGPYTH–) are present at the dimeric interface just like the corresponding active site motifs of DTD (– SQFTL– and –NXGPVT–) (**Figure 2A**). Besides the elements of DTD-like fold, specific interactions at the adenine-binding site for the recognition of A76 of tRNA are also highly conserved in ATD (**Figure 2—figure supplement 1**). Surprisingly, MmATD’s Gly-Pro motif occurs in *trans* conformation, unlike DTD’s Gly-Pro motif which always exists in *cis* conformation as observed in 107 protomers of 19 crystal structures from 5 different organisms (**Figure 2B,C**, **Figure 2—figure supplement 2A**). Notably, LmATD also possesses a cross-subunit Gly-*trans*Pro motif like MmATD (**Figure 2—figure supplement 2B**). Atomic B-factor analysis revealed that ATD’s Gly-*trans*Pro motif is rigid like DTD’s Gly-*cis*Pro motif (**Figure 2—figure supplement 3**). Additionally, ATD’s Gly-Pro residues exhibit a dramatic change of approximately 180° in ψ torsion angle when compared to DTD’s Gly-Pro residues due to remodeling of the local network of interactions in the vicinity of active site (**Figure 2D,E**, **Movie 1**). DTD’s Gly-*cis*Pro carbonyl oxygens are parallel and protrude into the active site pocket away from the protein core, *i.e.*, “outward parallel” orientation. ATD’s Gly-*trans*Pro carbonyl oxygens are also parallel, but they face away from the active site toward the protein core, *i.e.*, “inward parallel” orientation (**Figure 2B,C**). Thus, a direct consequence of *cis*-to-*trans* switch has a marked influence on the orientation of the carbonyl oxygens of glycine and proline residues of the Gly-Pro motif that is responsible for L-chiral rejection in DTD.

**Figure 2.**
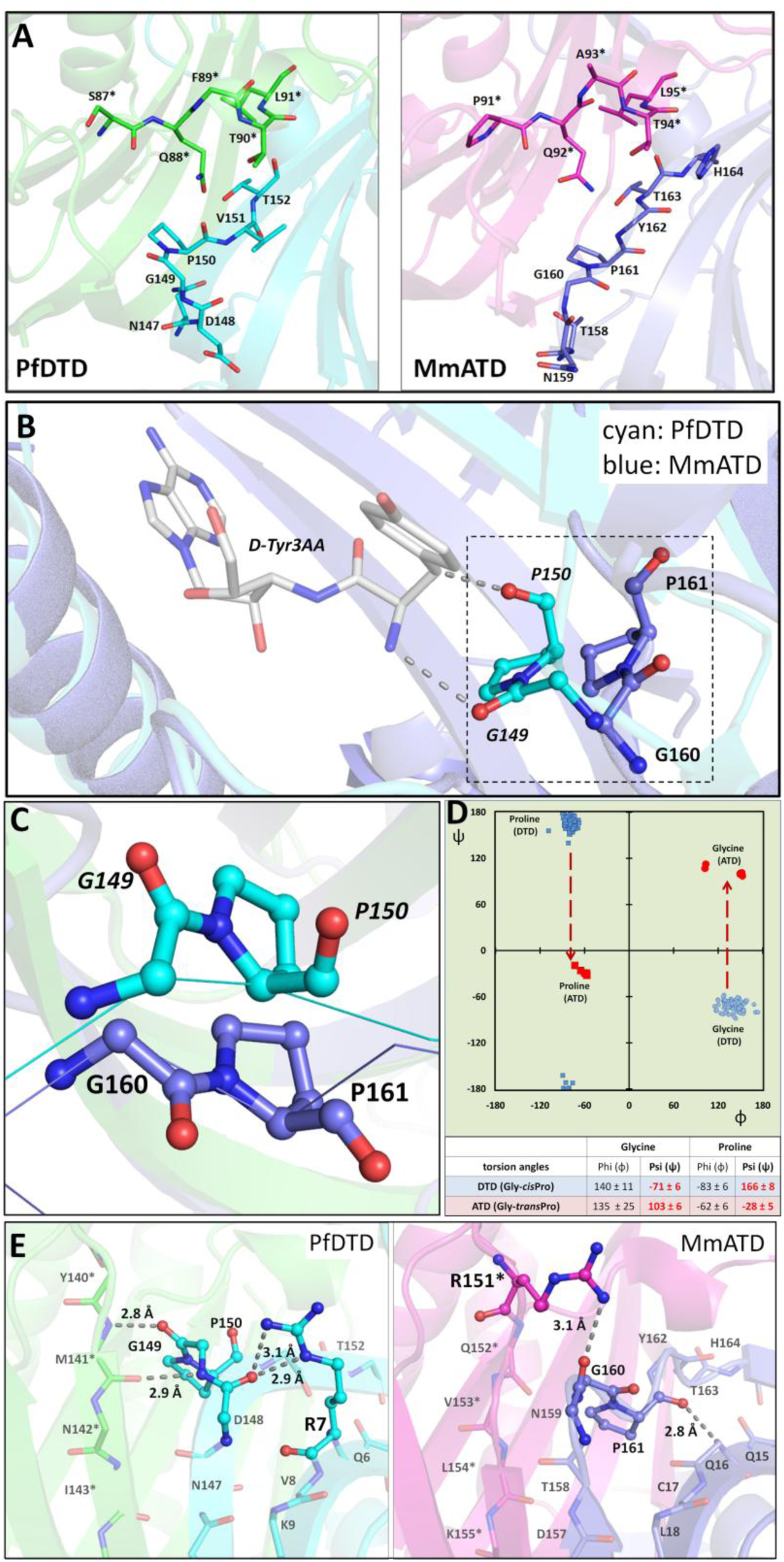
ATD has distinct active site elements/features as compared to DTD’s. (**A**) Crystal structures of PfDTD (PDB id: 4NBI) and MmATD showing that motifs 1 and 2 form the active site at the dimeric interface in both. (**B**) Comparison between Gly-*trans*Pro motif in MmATD and Gly-*cis*Pro motif in PfDTD (PDB id: 4NBI) after structural superposition of the two proteins. (**C**) The comparison shown in (**B**) depicted from a different angle, highlighting the opposite orientation of Gly-Pro carbonyl oxygens of the two proteins. (**D**) Ramachandran plot of glycine and proline residues of the Gly-Pro motif of all the available crystal structures of DTD (blue) and ATD (red), highlighting the change of ~180° in the ψ torsion angle. (**E**) Interaction of the side chain of Arg7 with the Gly-*cis*Pro motif of the same monomer in PfDTD (PDB id: 4NBI), and of the side chain of Arg151 with the Gly-*trans*Pro motif of the dimeric counterpart in MmATD. Residues from the dimeric counterpart are indicated by *.

### A conserved arginine near DTD’s active site has migrated in ATD

Upon further analysis of the active site region, it was observed that Arg7 in EcDTD or PfDTD, which is highly conserved in DTDs, is replaced by a conserved glutamine in ATD (Gln16 in MmATD). Interestingly, an invariant arginine is present in a totally different position in ATD (Arg151 in MmATD) (**Figure 1A**, **Figure 2E**). The side chain of Arg7 in PfDTD interacts with the main chain of Gly-*cis*Pro motif from the same monomer, thereby locking the motif rigidly in *cis* conformation (**Figure 2E**). This side chain–main chain interaction is conserved in all the available structures (107 protomers) of DTD (**Figure 2—figure supplement 4A**). In contrast, the interaction of MmATD’s Arg151 side chain with the main chain of Gly-*trans*Pro motif from the dimeric counterpart pulls the motif’s backbone outwards, thus holding the motif rigidly in *trans* conformation (**Figure 2E, figure supplement 4B**). Hence, the highly conserved arginine in the vicinity of DTD’s active site has migrated to a different position near ATD’s active site and is primarily responsible for the *cis*-to-*trans* switch of the Gly-Pro motif.

### ATD has “additional” space in its active site pocket due to flip of Gly-Pro carbonyl oxygens

In DTD, the “outward parallel” orientation of Gly-*cis*Pro carbonyl oxygens acts as a “chiral selectivity filter” to strictly reject all L-amino acids from the pocket through steric exclusion (**Ahmad et al., 2013; Routh et al., 2016**). Comparative analysis of active sites of DTD and ATD further revealed that the inward flip of ATD’s carbonyl oxygens due to *trans* conformation of its Gly-Pro motif has created “additional” space in its active site when compared to DTD. Consequently, ATD can easily accommodate a larger group in that space as opposed to just hydrogen in DTD (**Figure 2—figure supplement 5**, **Table S2**). This clearly suggests that ATD can cradle small L-amino acids in its active site pocket. The “additional” space created as a consequence of Gly-*trans*Pro switch in ATD prompted us to check its biochemical activity profile on L-aminoacyl-tRNAs, in addition to D-aminoacyl- and glycyl-tRNAs.

### ATD exhibits relaxed substrate chiral specificity due to *cis*-to-*trans* switch

The fact that ATD belongs to the DTD-like fold and its active site elements and architecture are similar to those of DTD indicated that it could be acting on some non-cognate aminoacyl-tRNA and hence could be involved in translational proofreading. Therefore, we generated the biochemical activity profile of MmATD by screening a spectrum of aminoacyl-tRNAs having either D-or L-amino acid as well as Gly-tRNA^Gly^. MmATD shows significant activity at 50 nM concentration on D-Tyr-tRNA^Tyr^, but fails to act on the L-counterpart even at 100-fold higher concentration (**Figure 3A,B**). It also deacylates Gly-tRNA^Gly^ at 500 nM concentration (**Figure 3C**). Thus, like DTD, ATD acts on both D-chiral and achiral substrates, albeit with significantly less efficiency. Strikingly, when tested with L-Ala-tRNA^Ala^, 500 nM MmATD displayed noticeable activity (**Figure 3D**). By contrast, EcDTD or PfDTD does not act on L-chiral substrates even at 100-fold higher concentration than required for D-chiral substrate (**Ahmad et al., 2013; Routh et al., 2016**). Therefore, biochemical probing suggested that ATD is an aminoacyl-tRNA deacylase with a relaxed specificity for substrate chirality, primarily due to the *trans* conformation of its active site Gly-Pro motif. It is for this reason that we named this protein ATD, which stands for <u>A</u>nimalia-specific <u>t</u>RNA <u>d</u>eacylase. ATD thus has a “gain of function” in L-chiral activity when compared to DTD due to the switch from Gly-*cis*Pro to Gly-*trans*Pro. Furthermore, biochemical data in conjunction with structural data indicate that ATD’s active site pocket, because of the “additional” space created by the inward movement of the Gly-Pro carbonyl oxygens, can accommodate only small L-amino acids like L-alanine but not the bulkier ones like L-tyrosine.

**Figure 3.**
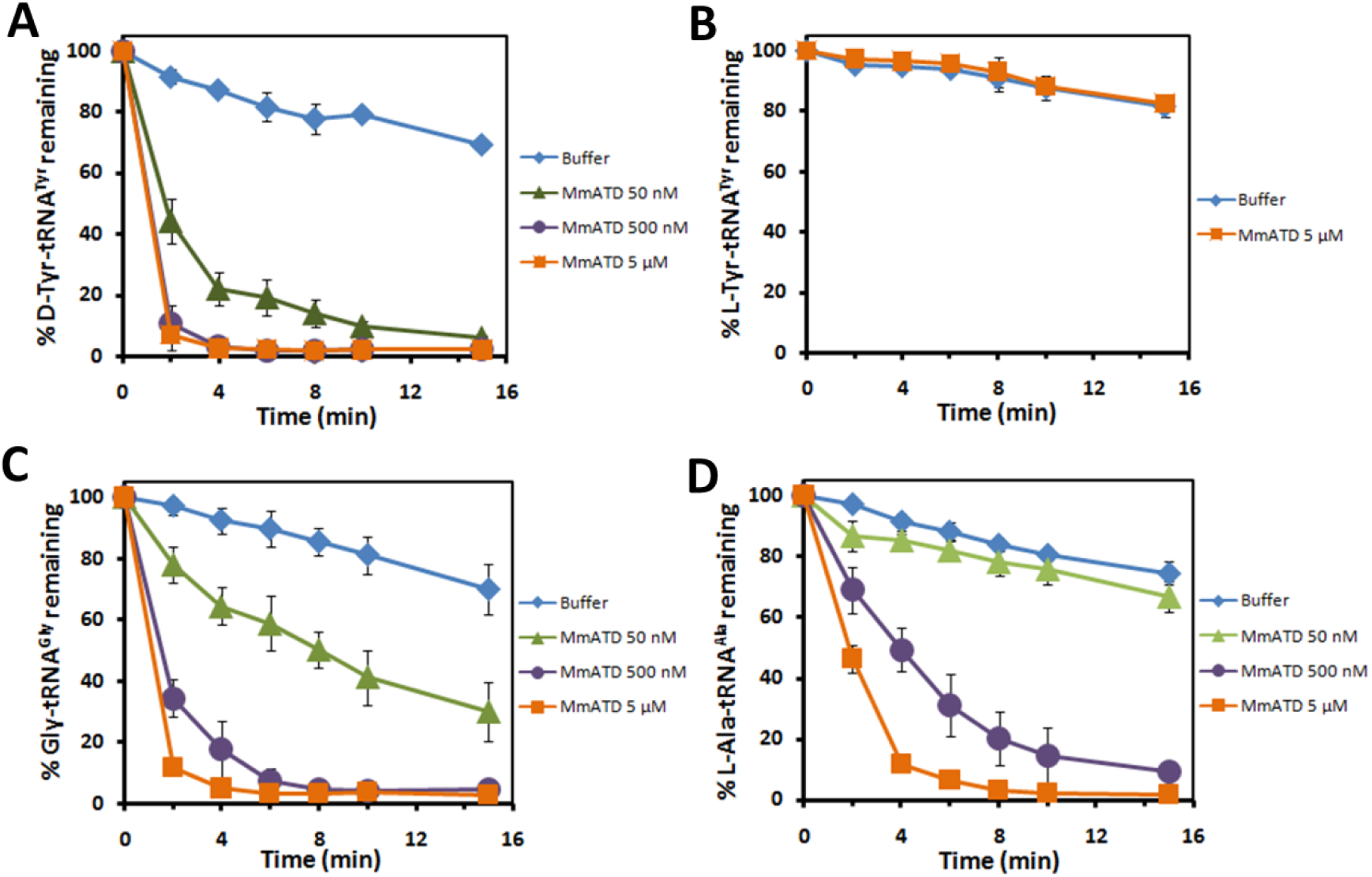
ATD displays relaxation of substrate chiral specificity. (**A**–**D**) Deacylation of DTyr-tRNA^Tyr^, L-Tyr-tRNA^Tyr^, Gly-tRNA^Gly^ and L-Ala-tRNA^Ala^ by different concentrations of MmATD. Error bars denote one standard deviation from the mean of three independent readings.

### ATD proofreads tRNA^Thr^(G4•U69) mischarged with L-alanine by eukaryotic AlaRS

While we were in the process of identifying the physiological substrate for ATD, a recent study reported that eukaryotic AlaRS has acquired the function of L-alanine mischarging on multiple non-cognate tRNAs harboring G4•U69 wobble base pair in the acceptor stem. One of the tRNAs that undergoes significant levels of such mischarging is tRNA^Thr^(G4•U69) (**Sun et al., 2016**). This is in addition to the canonical recognition of AlaRS-specific universally occurring G3•U70 in tRNA^Ala^ (**Hou and Schimmel, 1988; McClain and Foss, 1988**). Interestingly, such an error in the selection of tRNAs bearing G4•U69 was found only in the case of eukaryotic AlaRS and not the bacterial one (**Sun et al., 2016**). In this context, therefore, eukaryotic AlaRS can be called non-discriminating AlaRS (AlaRS^ND^), whereas bacterial AlaRS can be referred to as discriminating AlaRS (AlaRS^D^). The above finding prompted us to test the role of ATD in proofreading tRNA^Thr^(G4•U69) selection error made by eukaryotic AlaRS^ND^. Strikingly, biochemical assays with MmATD showed significantly higher selectivity for the non-cognate L-Ala-tRNA^Thr^(G4•U69) as the enzyme acts on the substrate at just 1 nM compared to its activity on the cognate L-Thr-tRNA^Thr^(G4•U69) at 50 nM (**Figure 4A**). Thus, MmATD displays a 50-fold difference in biochemical activity on these two substrates, indicating that L-Ala-tRNA^Thr^(G4•U69) is the preferred substrate for ATD.

**Figure 4.**
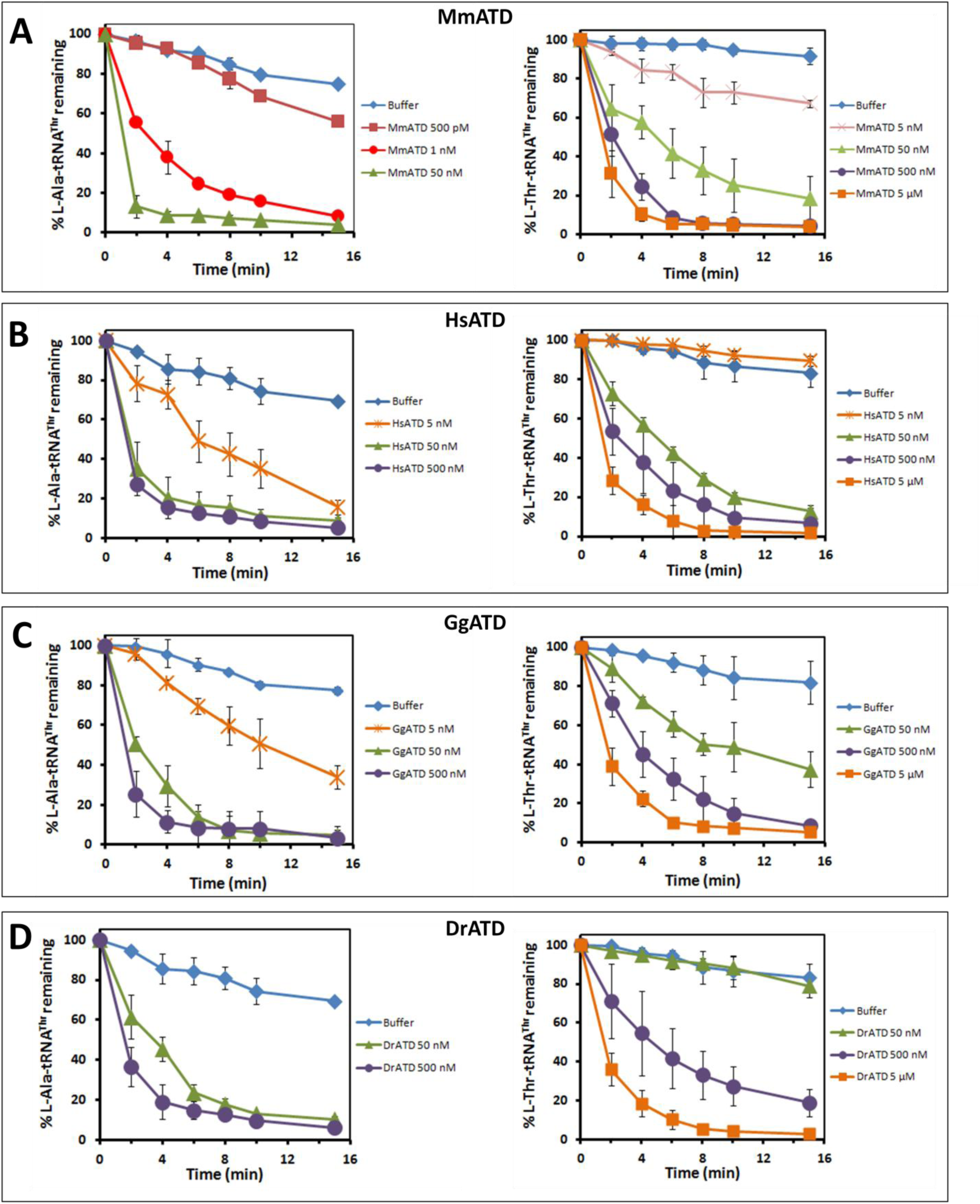
Proofreading of L-Ala-tRNA^Thr^(G4•U69) by ATD is conserved across organisms. Deacylation of D-Ala-tRNA^Thr^(G4•U69) and L-Thr-tRNA^Thr^(G4•U69) by different concentrations of (**A**) MmATD, (**B**) HsATD, (**C**) GgATD, and (**D**) DrATD. Error bars denote one standard deviation from the mean of three independent readings.

The other non-cognate tRNA that was found to be significantly mischarged by eukaryotic AlaRS was tRNA^Cys^(G4•U69) (**Sun et al., 2016**). We therefore checked ATD’s biochemical activity on L-Ala-tRNA^Cys^(G4•U69) to test its role in clearing the misacylated species. It was observed that MmATD acts on the substrate at 50 nM concentration (**Figure 4—figure supplement 1**). Thus, a comparison of biochemical activities of MmATD on different aminoacyl-tRNA substrates suggests that L-Ala-tRNA^Thr^(G4•U69) is the principal substrate of ATD, whereas L-Ala-tRNA^Cys^(G4•U69) may be partly cleared in the cellular context. The latter argument is supported by the observation that in the proteomics study of HeLa cells, misincorporation of L-alanine was observed at cysteine positions but not at threonine positions even though significant misacylation of tRNA^Thr^(G4•U69) with alanine was observed in *ex vivo* tRNA microarray experiments (**Sun et al., 2016**). Nevertheless, this observation is striking, since in humans, the enrichment of G4•U69 is significantly more in tRNA^Thr^ genes (20%) than in tRNA^Cys^ genes (3.4%) (**Table S3**).

### ATD’s biochemical activity is conserved in different organisms

To rule out any organism-specific phenomenon regarding ATD’s biochemical activity, we tested ATDs from multiple organisms belonging to different taxonomic groups under Chordata— human (*Homo sapiens*, HsATD) of class Mammalia (mammals), chicken (*Gallus gallus*, GgATD) of class Aves (birds), and zebrafish (*Danio rerio*, DrATD) of class Pisces (fishes). It was observed that all these ATDs can act on L-Ala-tRNA^Thr^(G4•U69) more efficiently than on L-Thr-tRNA^Thr^(G4•U69) (**Figure 4B–D**). Therefore, not only ATD’s activity on the non-cognate substrate but also its ability to discriminate between L-Ala-tRNA^Thr^(G4•U69) and L-Thr-tRNA^Thr^(G4•U69) is conserved across diverse classes of Chordata. We then analyzed tRNA^Thr^(G4•U69) gene sequences from diverse organisms which revealed that the first five base pairs in the acceptor stem are invariant or highly conserved (**Figure 4—figure supplement 2**). As ATD belongs to the DTD-like fold, its interaction with the tRNA is not expected to go beyond the first three or four base pairs in the acceptor stem. Hence, lack of variation in the acceptor stem of tRNA^Thr^(G4•U69) further suggests that ATD’s biochemical activity on L-Ala-tRNA^Thr^(G4•U69) is conserved across diverse taxonomic groups.

### EF-Tu protects the cognate L-Thr-tRNA^Thr^(G4•U69) but not the non-cognate L-Ala-tRNA^Thr^(G4•U69) from ATD

Since ATD had shown significant activity on L-Thr-tRNA^Thr^(G4•U69), we checked whether EF-Tu can protect the cognate substrate from ATD. EF-Tu occurs abundantly in the cell and most aminoacy L-tRNAs are expected to exist in complex with EF-Tu, ready for delivery to the ribosome (**Ishihama et al., 2008; Li et al., 2014**). On the basis of thermodynamic compensation, EF-Tu is expected to bind the non-cognate L-Ala-tRNA^Thr^(G4•U69) with significantly lower affinity compared to the cognate L-Thr-tRNA^Thr^(G4•U69) (**LaRiviere, 2001**). Competition assays demonstrated that L-Ala-tRNA^Thr^(G4•U69) undergoes significant deacylation with 50 nM MmATD in the presence of EF-Tu (**Figure 5A,B**). By contrast, L-Thr-tRNA^Thr^(G4•U69) is completely protected by EF-Tu even at 500 nM enzyme, whereas its protection is marginally relieved at 5 μM MmATD (**Figure 5C,D**). Hence, the discrimination potential/factor of M mATD for these two substrates enhances from approximately 50-fold in the absence of EF-Tu to more than 100-fold in the presence of EF-Tu (**Figure 4A**, **Figure 5B,D**). The above biochemical data clearly suggest that L-Ala-tRNA^Thr^(G4•U69) is ATD’s physiological substrate, and EF-Tu is able to confer adequate protection on the cognate substrate against ATD.

**Figure 5.**
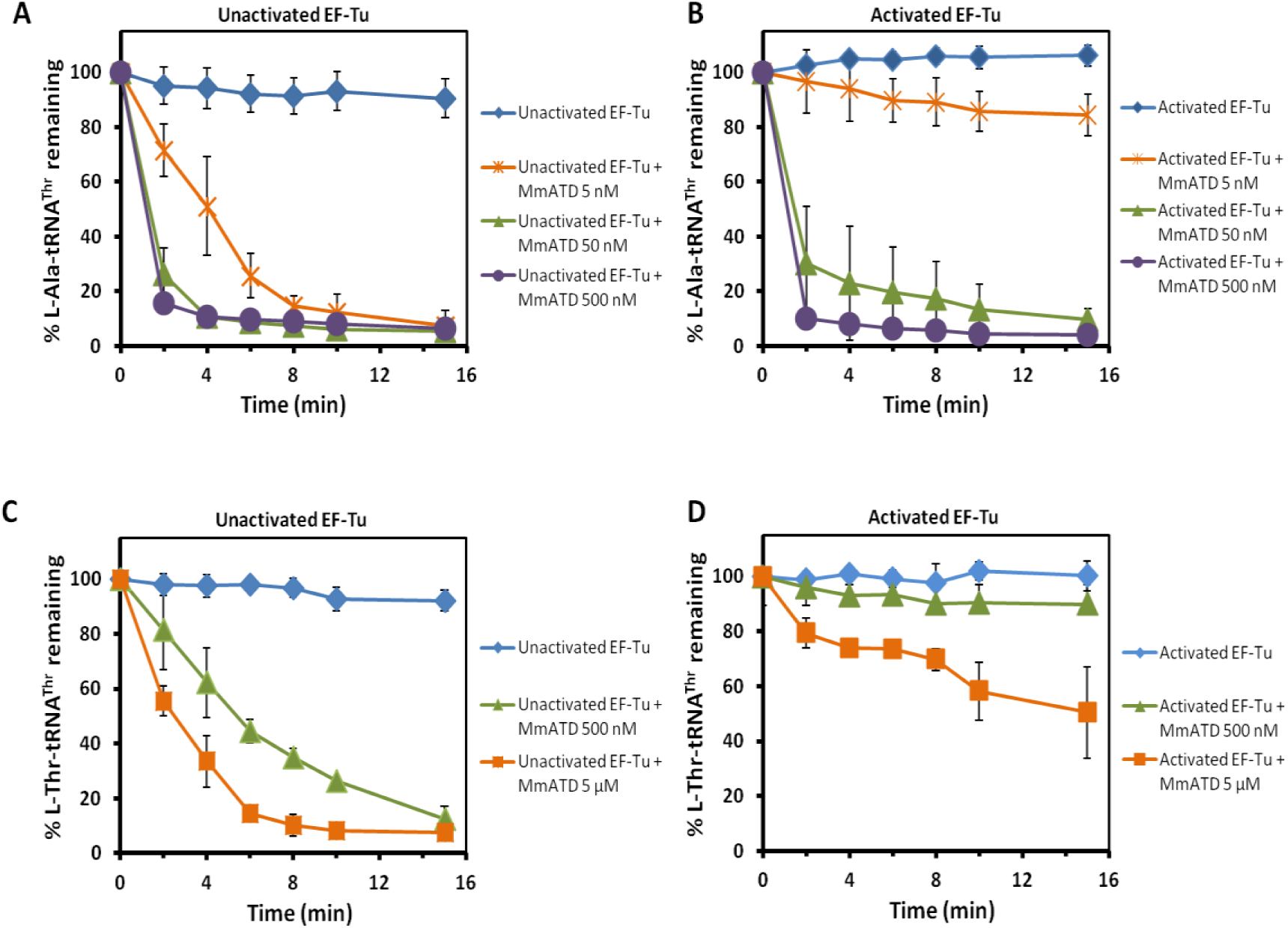
EF-Tu does not confer protection on the non-cognate L-Ala-tRNA^Thr^(G4•U69) against ATD. Deacylation of L-Ala-tRNA^Thr^(G4•U69) by different concentrations of MmATD in the presence of (**A**) unactivated EF-Tu, and (**B**) activated EF-Tu. Deacylation of L-ThrtRNA^Thr^(G4•U69) by different concentrations of MmATD in the presence of (**C**) unactivated EF-Tu, and (**D**) activated EF-Tu. Error bars denote one standard deviation from the mean of three independent readings.

### L-alanine mischarging on tRNA^Thr^(G4•U69) by AlaRS^ND^ and ATD have a strict and strong correlation

To ascertain whether any correlation exists between ATD and tRNA^Thr^(G4•U69), we performed a thorough bioinformatics analysis. It showed that many Animalia genomes are enriched in G4•U69-containing tRNA genes of which tRNA^Thr^(G4•U69) genes exhibit the highest level of enrichment. The enrichment of tRNA^Thr^(G4•U69) genes ranges from 20% to 40% (average ~30%). Such an enrichment of tRNA^Thr^(G4•U69) genes is observed throughout phylum Chordata as well as in one organism (*Strongylocentrotus purpuratus*) from phylum Echinodermata whose tRNA gene sequences are available in the database. The enrichment of G4•U69 is markedly less in other tRNA genes compared to tRNA^Thr^ genes. For example, tRNA^Cys^(G4•U69) genes constitute only 0.69–11.9% (average ~5%) of total tRNA^Cys^ genes (**Figure 6A,B**, **Table S3, Data 1**). Additionally, amongst all the G4•U69-containing tRNA genes found in Chordata, only tRNA^Thr^(G4•U69) genes occur in all those chordate species whose tRNA gene sequences are available in the database. Other tRNA genes carrying G4•U69 are restricted to only a small subset of organisms. For instance, tRNA^Cys^(G4•U69) genes occur in only 19 of 62 chordate species whose tRNA gene sequences are available in the database (**Figure 6C**). The above observations further indicate that L-Ala-tRNA^Thr^(G4•U69) constitutes the major substrate of ATD, whereas others including L-Ala-tRNA^Cys^(G4•U69) are only minor ones.

**Figure 6.**
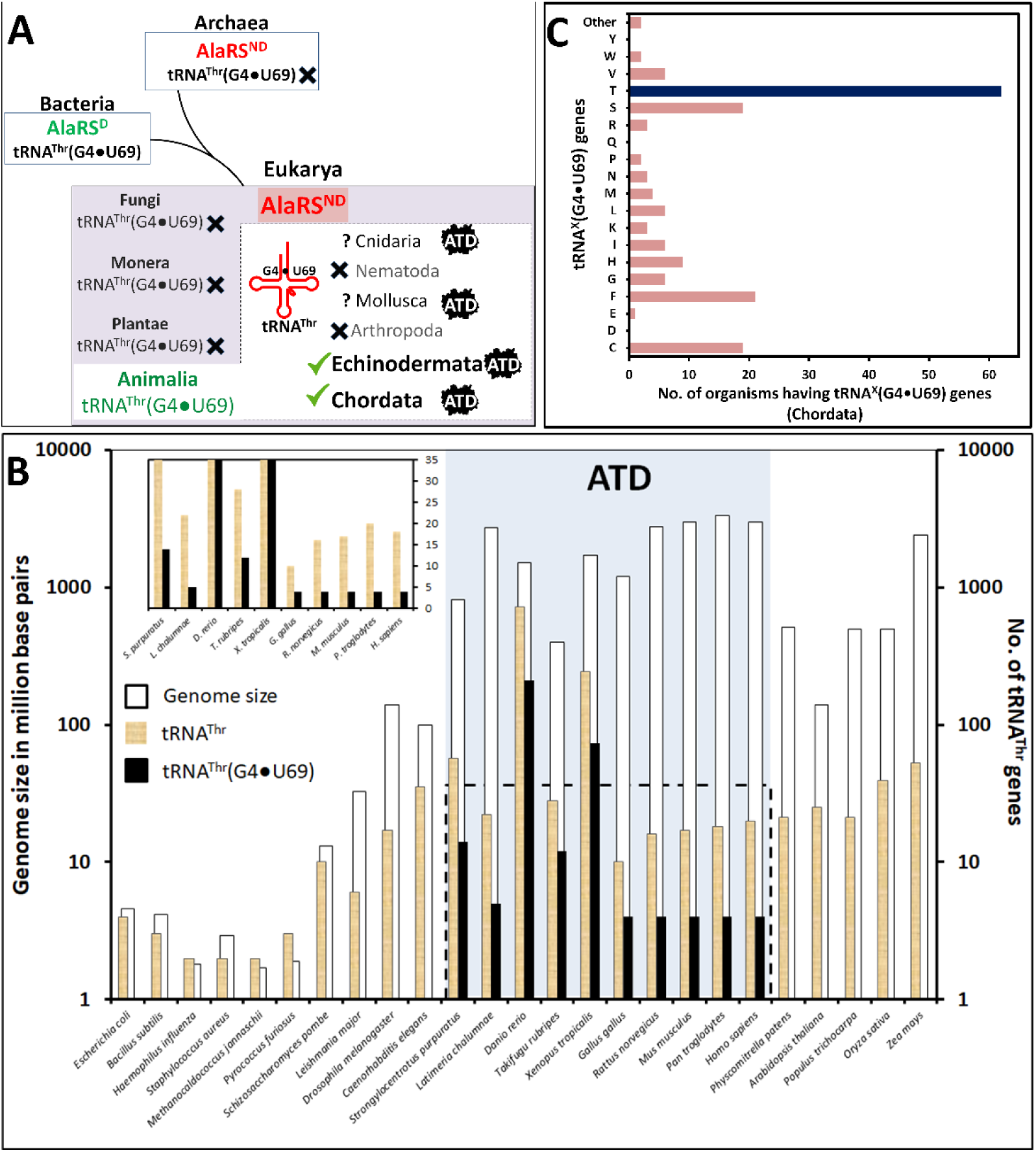
Enrichment of tRNA^Thr^(G4•U69) genes and presence of ATD show strict association. (**A**) Distribution of AlaRS^ND^, tRNA^Thr^(G4•U69) genes, and ATD in different domains of life. tRNA gene sequences of Cnidaria and Mollusca are not available in the database. (**B**) Bar graph (logarithmic scale) depicting genome size, total number of tRNA^Thr^ genes, and number of tRNA^Thr^(G4•U69) genes occurring in representative organisms belonging to all the three domains of life. Inset showing the number of total tRNA^Thr^ genes and tRNA^Thr^(G4•U69) genes in normal scale; genome size has not been shown for the sake of clarity. Presence of ATD is highlighted in light blue box. Data for occurrence of AlaRS^ND^ and tRNA^Thr^(G4•U69) genes have been taken from **Sun et al., 2016**. (**C**) Bar graph showing the number of organisms containing (G4•U69)-harboring tRNA genes which code for tRNAs specific for various proteinogenic amino acids.

Strikingly, a survey for the presence of ATD revealed its strict association with the enrichment of tRNA^Thr^(G4•U69) genes (**Figure 6A,B**). Remarkably, organisms (*e.g.*, *Drosophila melanogaster*) that lack tRNA^Thr^(G4•U69) genes simultaneously lack ATD, including archaea which also seem to possess eukaryotic-type AlaRS^ND^. Although many bacteria possess tRNA^Thr^(G4•U69) genes, they lack AlaRS^ND^ altogether and hence, the problem of L-alanine misacylation of tRNA^Thr^(G4•U69) does not arise at all. Thus, the problem of mischarging of tRNA^Thr^(G4•U69) with L-alanine arises only when tRNA^Thr^(G4•U69) is present alongside AlaRS^ND^. Such a strong as well as strict correlation between ATD and the problem of tRNA^Thr^(G4•U69) mis-selection by AlaRS^ND^, in terms of either concomitant occurrence or concomitant absence, clearly points toward a functional link between ATD and proofreading of error in tRNA^Thr^(G4•U69) selection.

## DISCUSSION

Overall, our extensive structural and biochemical probing in combination with in-depth *in silico* analysis confirms that ATD serves as a novel and dedicated factor for correcting the tRNA^Thr^(G4•U69) selection error committed by eukaryotic AlaRS^ND^ (**Figure 7**). The *trans* conformation of its active site Gly-Pro motif has led to a “gain of function” by relaxing its substrate chiral specificity. ATD thus rectifies a critical tRNA mis-selection rather than a mistake in amino acid selection by a synthetase that has been extensively studied so far in the context of proofreading (**Dock-Bregeon et al., 2000; Fersht, 1977, 1998; Fukai et al., 2000; Jakubowski and Fersht, 1981; Lincecum et al., 2003; Matinis and Boniecki, 2010; Nureki et al., 1998; Pauling, 1958; Perona and Gruic-Sovulj, 2014; Silvian et al., 1999; Yadavalli and Ibba, 2012**). Such an error correction capability has not been attributed to any of the known editing domains, although ambiguous tRNA selection happens in several instances, wherein the ambiguity imparts selective advantage to the system (**Figure 7**) (**Schwartz and Pan, 2017; Shepherd and Ibba, 2014**). Besides, it also suggests that the evolutionary gain of function by AlaRS^ND^ in charging G4•U69-bearing tRNAs with L-alanine may be advantageous, but may also require factors like ATD to keep such “errors” below precarious levels, thus avoiding global misfolding and cell death.

**Figure 7.**
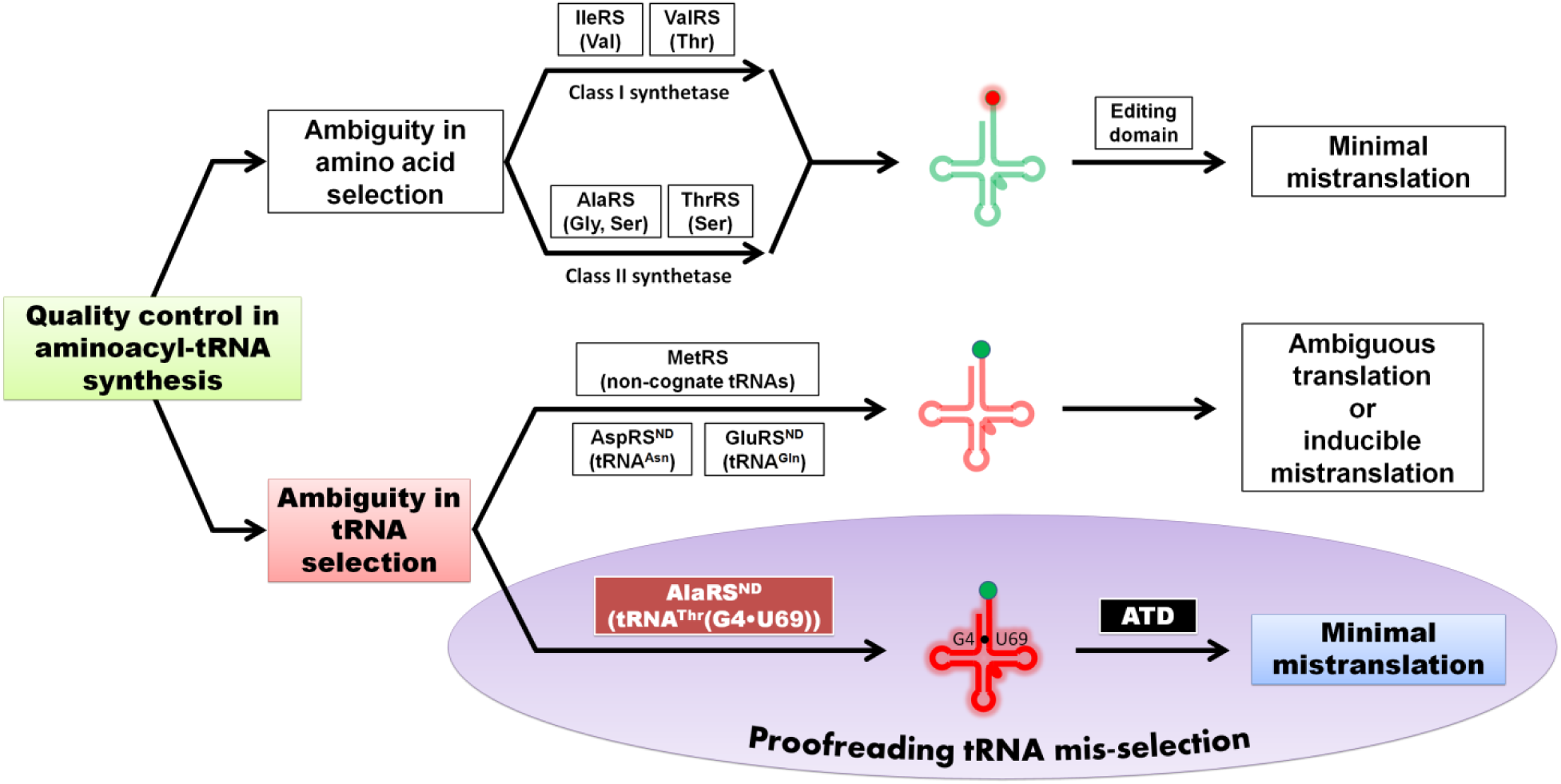
ATD is a unique and dedicated proofreading factor that rectifies a critical tRNA selection error. Model for mis-selection and consequent misacylation of tRNA^Thr^(G4•U69) with L-alanine by AlaRS^ND^, and its subsequent proofreading by ATD. Cognate and non-cognate tRNAs (clover leaf model) are colored in green and red, respectively. Likewise, cognate and non-cognate amino acids (circle) are rendered in green and red, respectively.

The role of ATD in Animalia to specifically prevent, minimize or regulate substitution of L-alanine for L-threonine in proteins may be crucial. It is tempting to speculate that threonine-to-alanine mutations will modulate a diverse array of phosphorylation sites on proteins, thereby causing a drastic modification of the cellular phosphoproteome. The regulatory function, if any, of ATD in generating such proteome diversity, thereby providing selective advantage to a cell or tissue type and under specific conditions such as pathogenesis or immune response, needs to be explored through up/down-regulation as well as by using knockout approaches in multiple systems. It has been recently suggested that the size of tRNA limits the identity determinants required for faithful translation without cross-reacting with non-cognate synthetases (**Saint-Léger et al., 2016**). As has been noted in the current work, the expansion of genome size (from around 100 million base pairs in non-Chordata to >1 billion base pairs in Chordata) has led to such a cross-reactivity and enhancement in tRNA mis-selection, thereby necessitating recruitment of dedicated factors for error correction (**Figure 6B**). Identification of ATD in the present study provides the first instance of such a scenario. The advent of ATD thus marks a key event associated with the appearance of Animalia, and more specifically of Chordata about 500 million years ago, to ensure translational quality control.

## Materials and Methods

### Cloning, expression and purification

The genes encoding ATDs and *M. musculus* ThrRS(ΔNTD) (residue 320–721) were PCR-amplified from respective cDNAs and inserted into pET-28b vector using restriction-free cloning (**van den Ent and Löwe, 2006**). To generate C-terminal 6X His-tagged protein, the stop codon was removed from the reverse primer. *M. musculus* full-length AlaRS gene was PCR-amplified from mouse cDNA and inserted into pET-28b vector between *Nde*I and *Xho*I sites using conventional restriction-based cloning. The recombinant proteins were overexpressed in Rosetta^TM^ (DE3) strain of *E. coli*. Purification of His-tagged proteins (ATDs, AlaRS and ThrRS) was done using a two-step protocol, *i.e.*, Ni-NTA affinity chromatography followed by size exclusion chromatography (SEC). The storage buffer for ATDs contained 100 mM Tris (pH 8.0) and 200 mM NaCl, while that for AlaRS and ThrRS comprised 50 mM Tris (pH 8.0), 150 mM NaCl and 5 mM 2-mercaptoethanol (β-ME). Un-tagged MmATD was purified using cation exchange chromatography (CEC) followed by SEC. For CEC, ATD-overexpressed *E. coli* cell pellet was lysed in a buffer containing 50 mM Bis-Tris (pH 6.5) and 20 mM NaCl, and the supernatant was subjected to chromatographic separation using sulfopropyl sepharose (GE Healthcare Life Sciences, USA). The protein of interest was eluted using a NaCl gradient of 20 mM to 500 mM. For SEC, Superdex 75 column (GE Healthcare Life Sciences, USA) was used.

### Crystallization of MmATD

The purified un-tagged MmATD was screened for crystallization conditions using different screens (Index, Crystal Screen HT, PEG/Ion and PEGRx from Hampton Research, USA) at two different temperatures—4°C and 20°C. Mosquito Crystal (TTP LabTech, UK) crystallization robot was used to set up crystallization experiments using sitting-drop vapor diffusion method by mixing 1 μl protein and 1 μl reservoir buffer in a 96-well MRC plate with three sub-wells (Molecular Dimensions, UK). The initial hits from the screens were further expanded for optimization using sitting-drop vapor diffusion method in 96-well format MRC plates having three sub-wells. Reservoir buffer with 0.1 M Bicine (pH 8.0) and 15% PEG1500 yielded good diffraction-quality crystals.

### X-ray diffraction data collection, structure solution and refinement

In-house X-ray facility consisting of RigakuMicromax007 HF with rotating-anode generator and MAR345-dtb image plate detector was used for crystal screening and data collection at 100 K using an Oxford Cryostreamcooler (Oxford Cryosystems, UK). The wavelength of X-rays used was 1.5418 Å, corresponding to Cu Kα radiation. HKL2000 (**Otwinowski and Minor, 1997**) was used for data processing and MOLREP-AUTO MR from CCP4 suite (**CCP4, 1994**) for molecular replacement. PfDTD (PDB id: 4NBI), with the ligand removed, was used as the search model for molecular replacement. Refinement and model building were done using REFMAC (**Murshudov et al., 1997**) and COOT (**Emsley and Cowtan, 2004**), respectively. Structure validation was done using PROCHECK (**Laskowski et al., 1993**). PyMOL Molecular Graphics System, Version 1.7.6.0 Schrödinger, LLC was used to generate figures. Structure-based multiple sequence alignment was carried out using the T-Coffee server in Expresso mode (http://tcoffee.crg.cat/apps/tcoffee/do:expresso), and the corresponding figure was generated using ESPript 3.0 (http://espript.ibcp.fr/ESPript/cgi-bin/ESPript.cgi). The atomic coordinates of MmATD crystal structure have been deposited in the Protein Data Bank with accession code 5XAQ.

### *In vitro* biochemical experiments

tRNAs (*M. musculus* tRNA^Gly^, tRNA^Ala^, tRNA^Thr^(G4•U69) and *E. coli* tRNA^Tyr^) were generated by *in vitro* transcription of the corresponding tRNA genes using MEGAshortscript T7 Transcription Kit (Thermo Fisher Scientific, USA). tRNAs were end-labeled with [α-^32^P] ATP (BRIT-Jonaki, India) using CCA-adding enzyme (**Ledoux and Uhlenbeck, 2008**). Glycylation of tRNA^Gly^ was done by incubating 1 μM tRNA^Gly^ with 2 μM *Thermus thermophilus* GlyRS in a buffer containing 100 mM HEPES (pH 7.5), 10 mM KCl, 30 mM MgCl2, 50 mM glycine and 2 mM ATP at 37°C for 15 min. Alanylation of tRNA^Ala^ and tRNA^Thr^(G4•U69) was performed by incubating 1 μM tRNA^Ala^ and 10 μM tRNA^Thr^(G4•U69) with 8 μM *M. musculus* AlaRS in a solution composed of 100 mM HEPES (pH 7.5), 30 mM KCl, 100 mM MgCl2, 10 mM ATP, 10 mM dithiothreitol (DTT), 1 unit/ml of PPase enzyme (Thermo Fisher Scientific, USA) and 100 mM L-alanine at 37°C for 15 min. Threonylation of tRNA^Thr^(G4•U69) was carried out by incubating 1 μM tRNA^Thr^(G4•U69) and *M. musculus* ThrRS(ΔNTD) in a buffer comprising 100 mM HEPES (pH 7.5), 100 mM MgCl_2_, 300 mM KCl, 45 mM L-threonine, 2.5 mM DTT and 2 mM ATP at 37°C for 15 min. tRNA^Tyr^ was aminoacylated by *E. coli* TyrRS as described in **Ahmad et al., 2013**. *T. thermophilus* EF-Tu activation and protection assays were done using the protocol as described in **Routh et al., 2016**. Deacylation assays were carried out in conditions as described in **Ahmad et al., 2013** and EF-Tu rescue experiments as described in **Routh et al., 2016**. All the experiments were performed in triplicates, and the mean values were used to plot the graphs. Error bars denote one standard deviation from the mean of three independent readings.

### Bioinformatic analysis

Protein sequences were retrieved from NCBI and were subjected to phylogenetic tree construction using the web server http://www.phylogeny.fr/ (bootstrap number = 100) and iTOL web server http://itol.embl.de/. tRNA gene sequences were retrieved from GtRNAdb and sequences having tRNAscan-SE score >50 were used for analysis (http://gtrnadb.ucsc.edu/). The list of organisms whose genomes have been completely sequenced was obtained from KEGG GENOME database (http://www.genome.jp/kegg/genome.html). Information about genome size was taken from the web server http://www.bionumbers.hms.harvard.edu/default.aspx. Multiple sequence alignment of tRNA^Thr^ and tRNA^Thr^(G4•U69) was prepared using T-Coffee server in M-Coffee mode (http://tcoffee.crg.cat/apps/tcoffee/do:mcoffee), while consensus sequence logo was prepared using WebLogo server (http://weblogo.berkeley.edu/logo.cgi).

### Movie preparation

The two conformations/states, one of PfDTD (initial state; PDB id: 4NBI) and the other of MmATD, were morphed using UCSF Chimera software (**Pettersen et al., 2004**). Movie was then prepared using PyMOL Molecular Graphics System, Version 1.7.6.0 Schrödinger, LLC.

## Acknowledgements

The authors acknowledge Dr. P. Chandra Shekar for kindly providing mouse cDNA. S.K.K. thanks DST-INSPIRE, India, for research fellowship. M.M. thanks Department of Biotechnology, India for research fellowship. S.B.R. and R.S. acknowledge funding from 12th Five Year Plan Project BSC0113 of CSIR, India. R.S. also acknowledges funding from J.C. Bose Fellowship of SERB, India, and Centre of Excellence Project of Department of Biotechnology, India.

## Supplementary figures and tables

**Figure 1—figure supplement 1.**
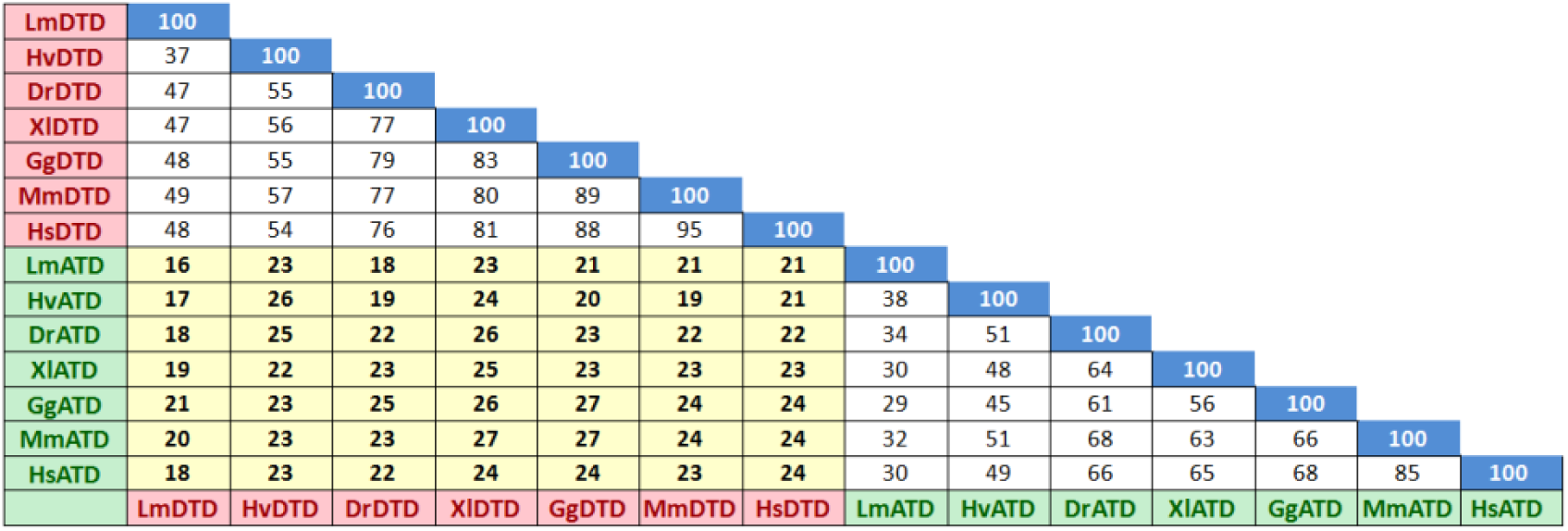
DTD and ATD show less than 30% sequence identity. Matrix showing percentage identities between DTDs, between ATDs as well as between DTDs and ATDs belonging to representative organisms. Lm, *Leishmania major*; Hv, *Hydra vulgaris*; Dr, *Danio rerio*; Xl, *Xenopus laevis*; Gg, *Gallus gallus*; Mm, *Mus musculus*; Hs, *Homo sapiens*.

**Figure 1—figure supplement 2.**
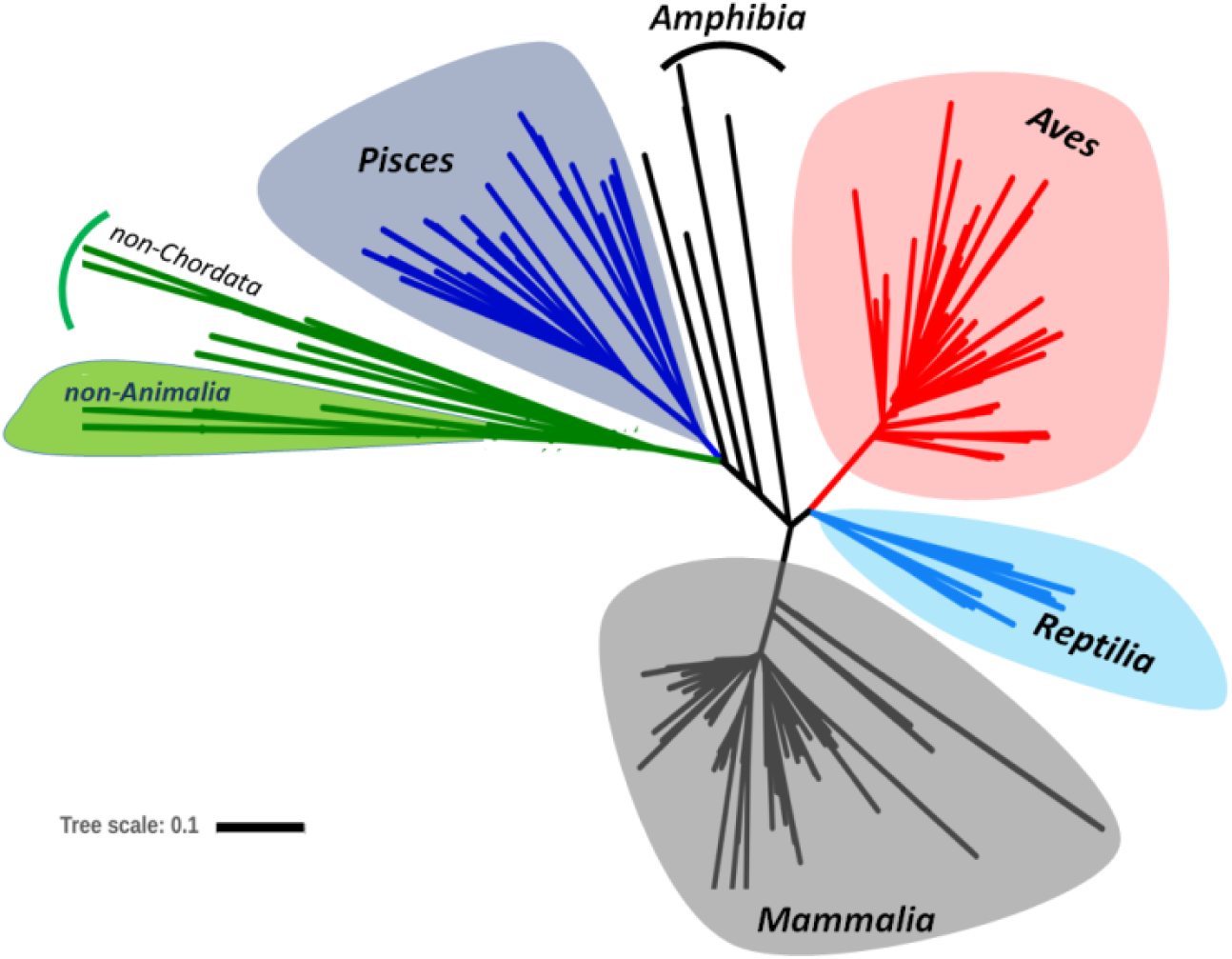
ATD exhibits exclusive distribution in kingdom Animalia. Phylogenetic analysis depicting the presence of ATD mainly in kingdom Animalia, and more specifically in phylum Chordata, which comprises Pisces, Amphibia, Reptilia, Aves and Mammalia. The few non-Animalia that harbor ATD are mostly parasites of various chordates, hence the possibility of horizontal transfer of ATD gene.

**Figure 1—figure supplement 3.**
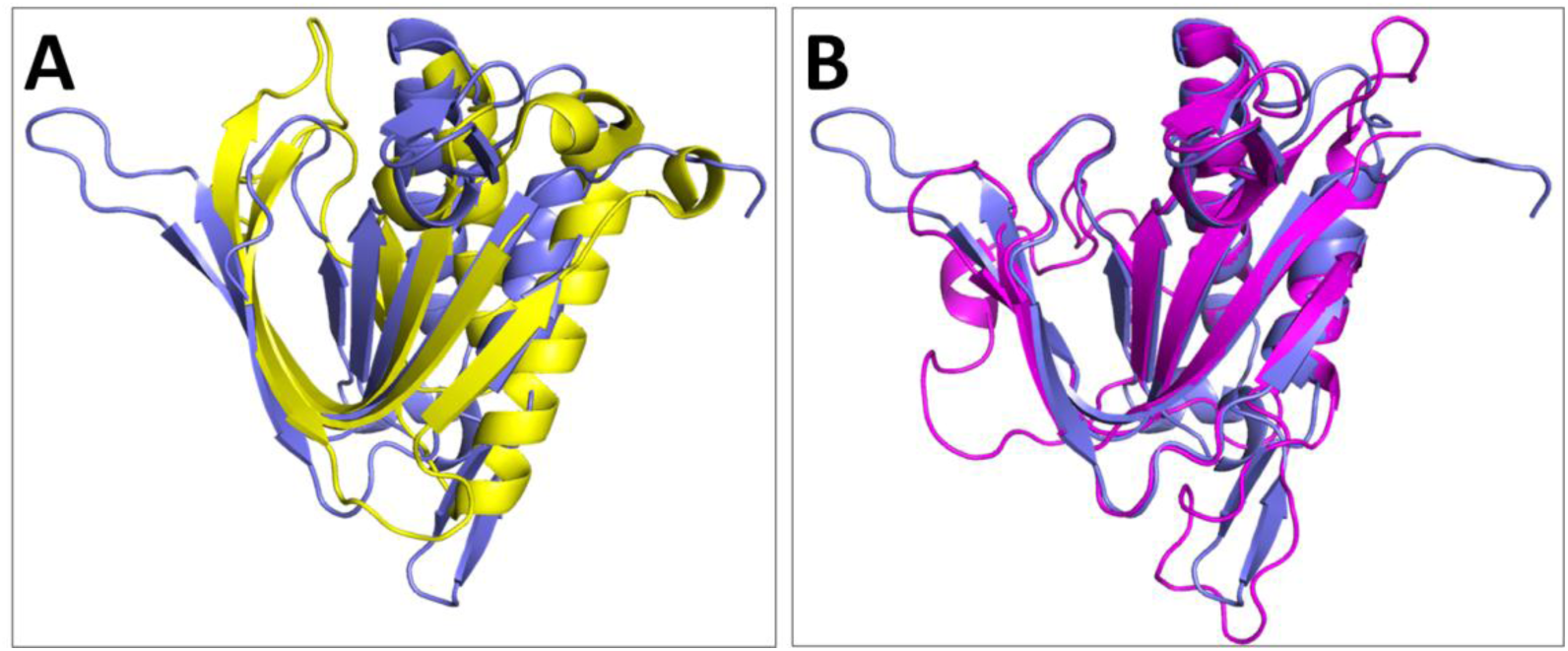
ATD and NTD are structural homologs belonging to the DTD-like fold. (**A**) Structural overlap of MmATD monomer (blue) on PabNTD monomer (yellow; PDB id: 3PD3) (r.m.s.d., 3.34 Å over 77 Cα atoms) showing that the two belong to the same fold. (**B**) Structural overlap of MmATD monomer (blue) on LmATD monomer (magenta; PDB id: 1TC5) (r.m.s.d., 1.29 Å over 148 Cα atoms).

**Figure 2—figure supplement 1.**
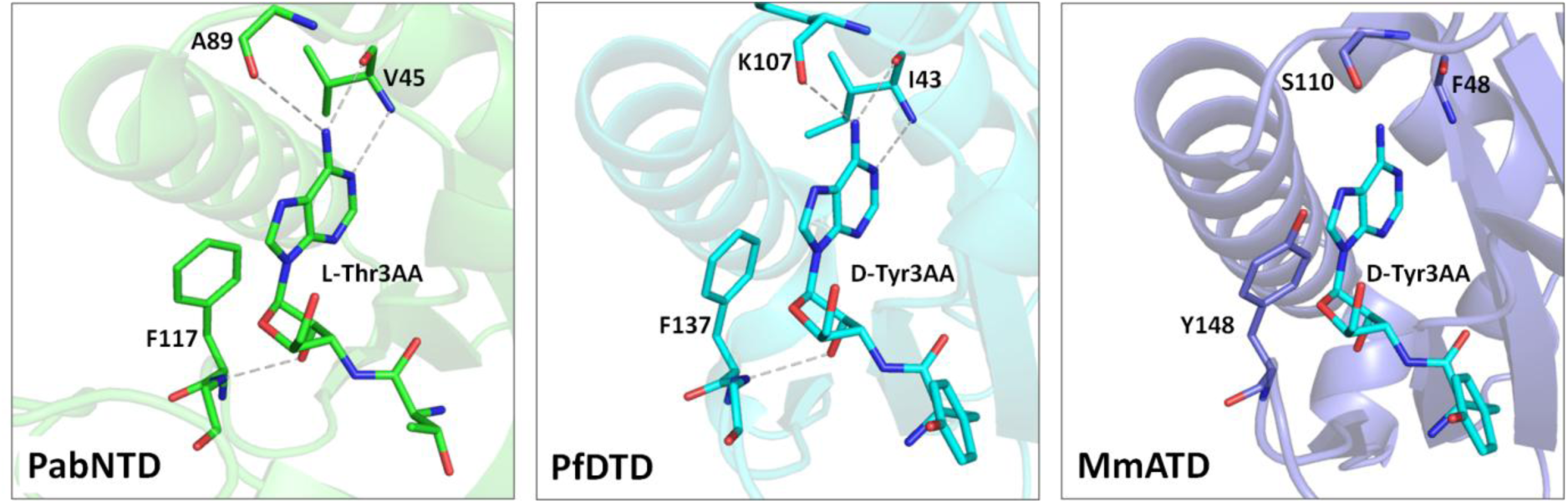
The adenine-binding pocket is conserved in the DTD-like fold. The elements of adenine-binding pocket of the DTD-like fold, encompassing NTD, DTD and ATD, are conserved. The ligand D-tyrosyl-3′-aminoadenosine (D-Tyr3AA) in MmATD has been modeled on the basis of structural overlap of MmATD on PfDTD (PDB id: 4NBI). The PDB id for PabNTD is 3PD3.

**Figure 2—figure supplement 2.**
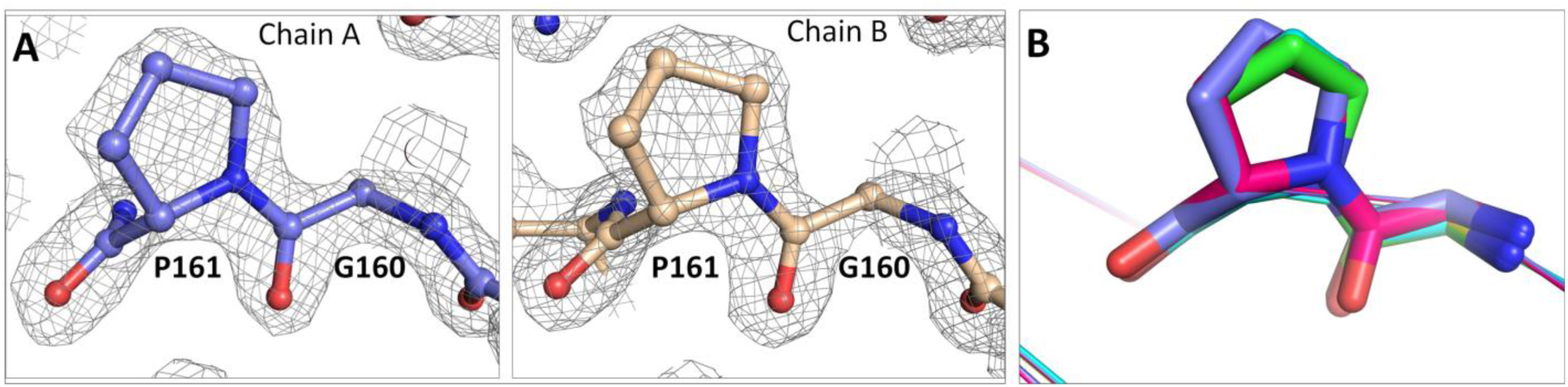
ATD has an active site Gly-*trans*Pro motif. (**A**) (2*F_o_-F_c_*) map, contoured at 2σ, showing clean density for the Gly-*trans*Pro motif of both protomers present in the asymmetric unit of MmATD crystal structure. (**B**) Structural superposition of LmATD protomers (cyan, green, purple and yellow; PDB id: 1TC5) on MmATD protomers (blue and magenta) highlighting the *trans* conformation of the active site Gly-Pro motif in both proteins.

**Figure 2—figure supplement 3.**
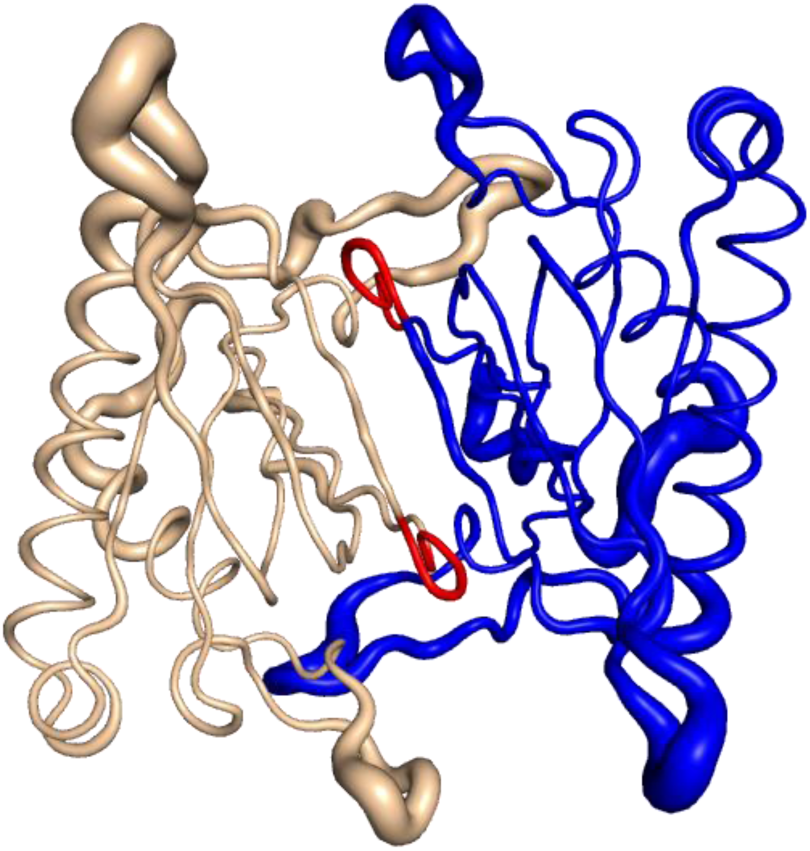
Atomic B-factor analysis of MmATD shows rigidity of Gly-*trans*Pro motif. Backbone representation of MmATD showing the variation in atomic B-factor. Regions represented as thin lines are more rigid and therefore have low B-factor values, whereas those depicted as thick lines are more flexible and thus have higher values. The atomic B-factor for the structure lies in the range 18–62 Å^2^. Regions depicted in red represent glycine and proline residues of the Gly-*trans*Pro motif, whose average B-factor is 24 Å ^2^ (B-factor of protein is 31 Å^2^). This value is similar to that of PfDTD (B-factors of Gly-*cis*Pro motif and protein are 25 Å^2^ and 29 Å^2^, respectively; PDB id: 4NBI), showing that the Gly-*trans*Pro motif in ATD is rigidly fixed like the Gly-*cis*Pro motif in DTD. The two monomers of ATD homodimer have been rendered in different colours.

**Figure 2—figure supplement 4.**
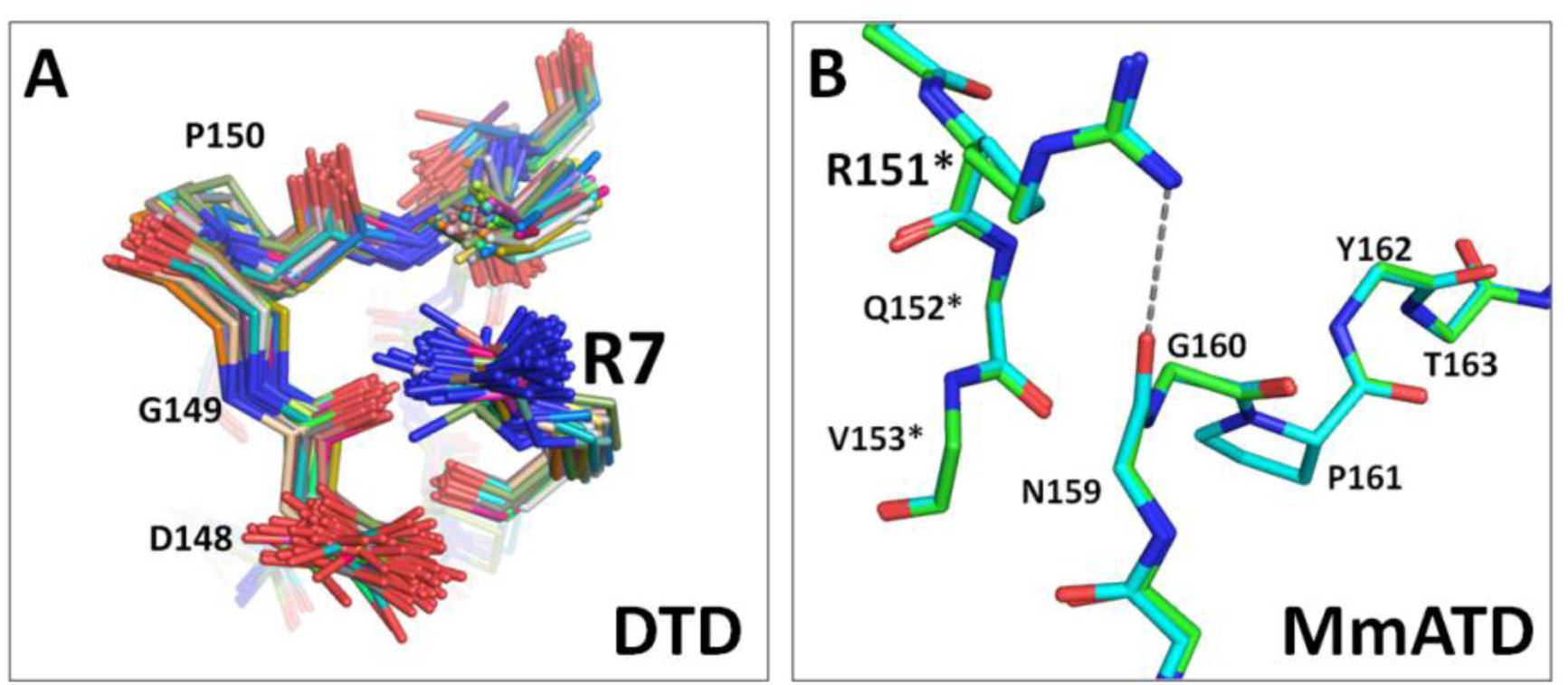
An arginine is involved in the rigid fixation of Gly-Pro motifs in DTD and ATD. (**A**) Structural superposition of 107 protomers of DTD from 19 PDBs and 5 different organisms showing rigid fixation of Gly-*cis*Pro motif by a conserved interaction with the side chain of a highly conserved arginine (Arg7, PfDTD) from the same monomer. Numbering of residues is according to PfDTD. (**B**) Structural overlap of the two protomers of MmATD showing that Gly-*trans*Pro motif in ATD is firmly fixed by a conserved interaction with the side chain of an invariant arginine (Arg151). Residues from the dimeric counterpart are indicated by *.

**Figure 2—figure supplement 5.**
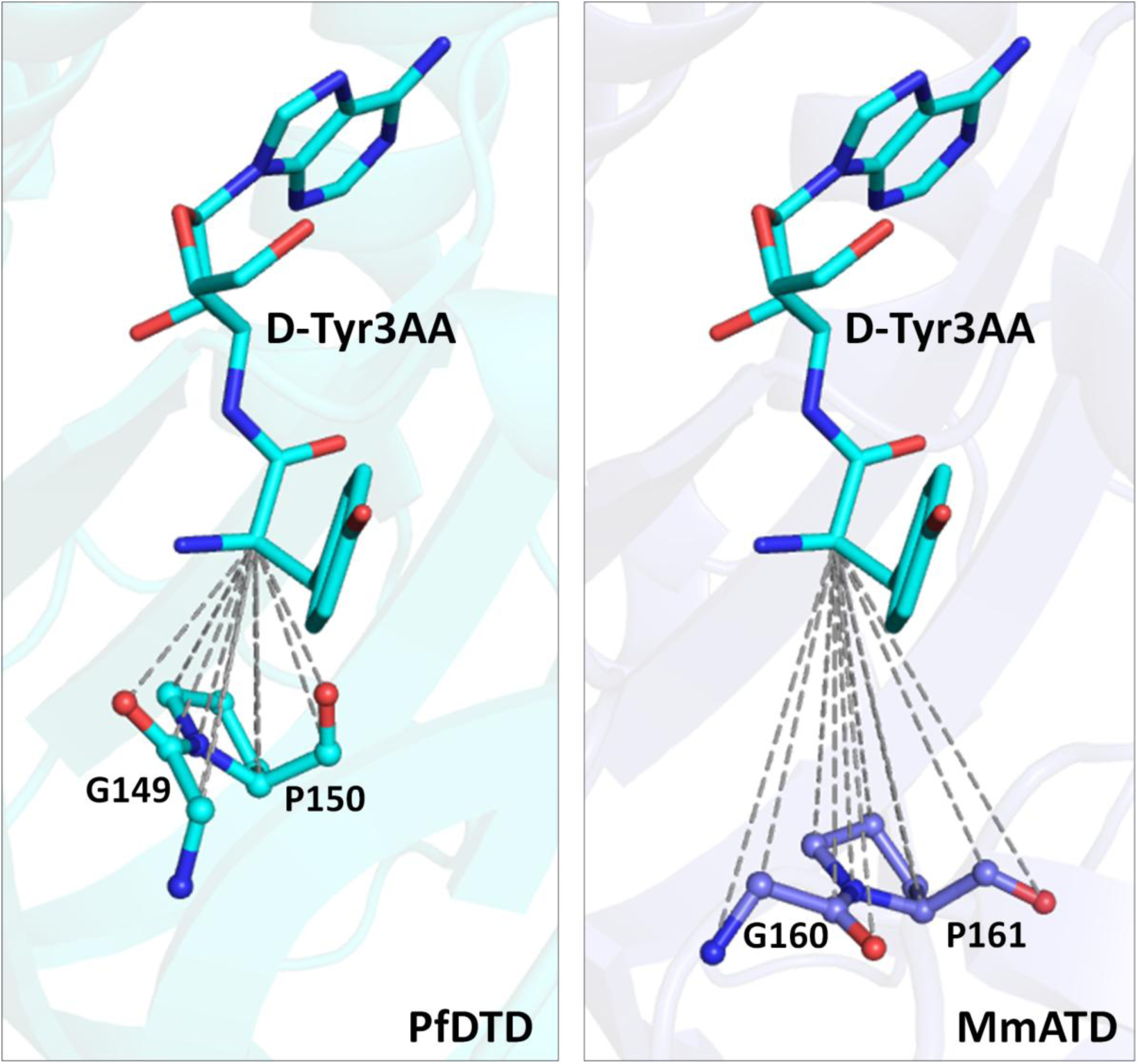
ATD has “additional” space in its active site pocket compared to DTD. Comparison between active site pockets of PfDTD (PDB id: 4NBI) and MmATD showing “additional” space in the latter due to the inward movement of Gly-Pro carbonyl oxygens. For MmATD, the ligand was modeled in the active site after superposition of MmATD dimer on PfDTD dimer.

**Figure 4—figure supplement 1.**
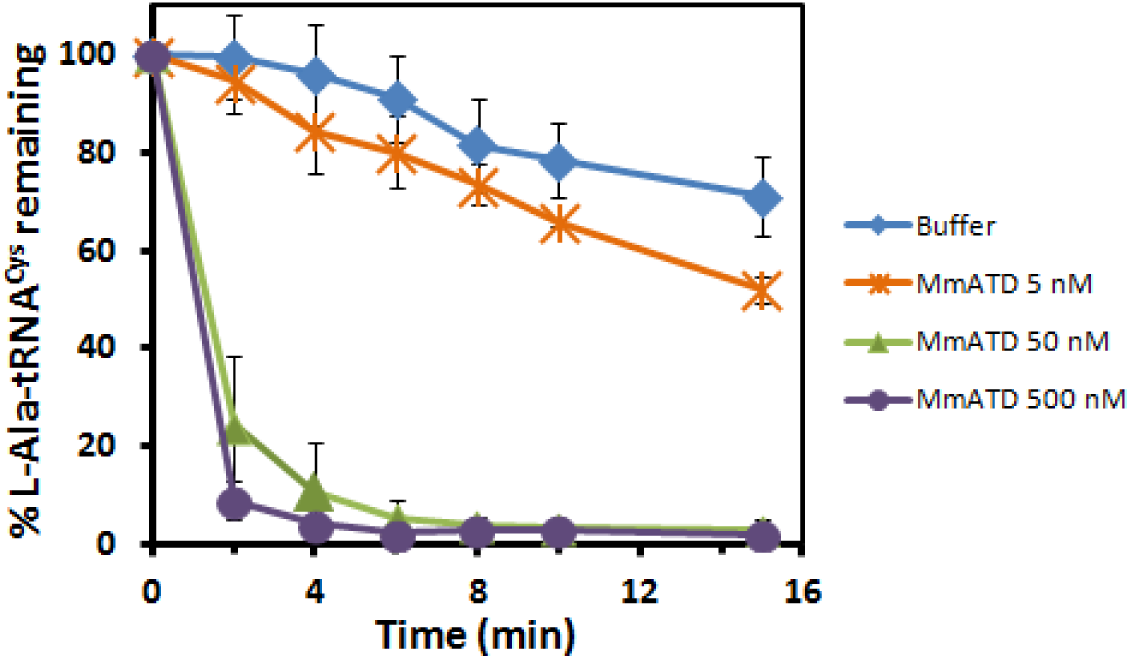
ATD can also act on L-Ala-tRNA^Cys^(G4•U69). Deacylation of L-Ala-tRNA^Cys^(G4•U69) by different concentrations of MmATD.

**Figure 4—figure supplement 2.**
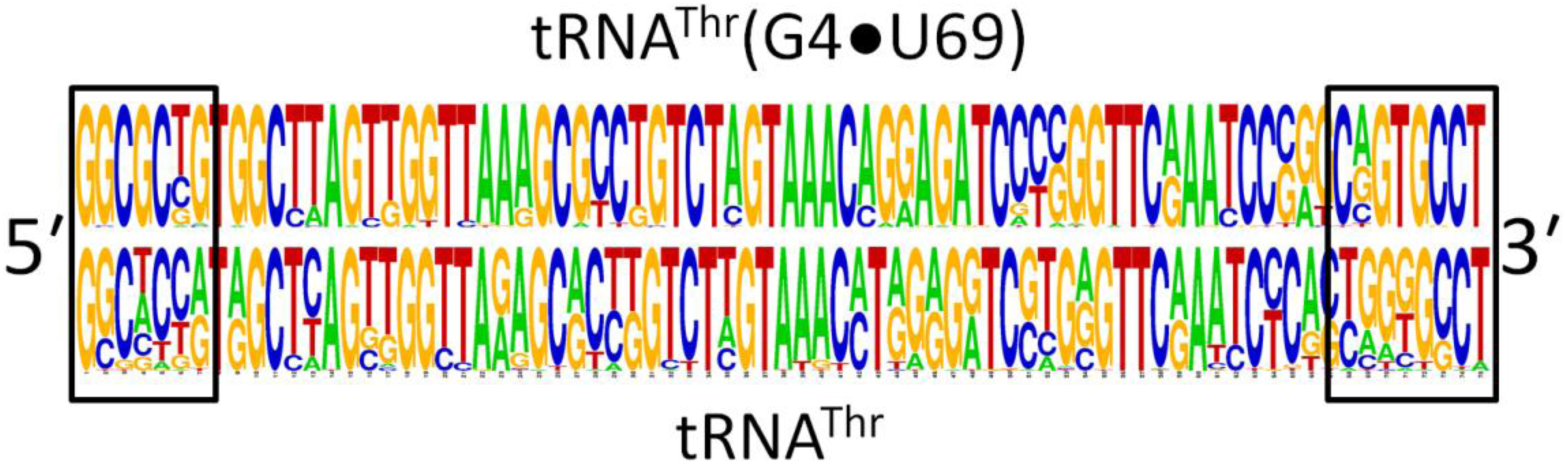
Acceptor stem elements of tRNA^Thr^(G4•U69) genes are highly conserved. Consensus sequence showing significantly higher conservation of acceptor stem residues (enclosed in boxes) in tRNA^Thr^(G4•U69) genes than in non-G4•U69-containing tRNA^Thr^ genes. The gene sequences taken for analysis belong to the following representative organisms: *Strongylocentrotus purpuratus*, *Latimeria chalumnae*, *G. gallus*, *M. musculus*, *H. sapiens*.

**Table S1.** Crystallographic data collection and refinement statistics

| **MmATD** |  |
| --- | --- |
| **Data Collection** |  |
| Space group | $P2_12_12_1$ |
| Cell dimensions: |  |
| $a$ (Å) | 43.37 |
| $b$ (Å) | 75.35 |
| $c$ (Å) | 103.61 |
| Resolution range (Å)* | 50–1.86 (1.93–1.86) |
| Total Observations | 166914 |
| Unique reflections | 29268 (1473) |
| Completeness (%) | 89.5 (51.2) |
| $R_{\text{merge}}$ (%) | 7.6 (38.0) |
| $\langle I/(\sigma)I \rangle$ | 21.8 (2.4) |
| Redundancy | 6.4 (3.7) |
| **Data refinement** |  |
| Resolution (Å) | 1.86 |
| No. of reflections | 24840 |
| $R_{\text{work}}$ (%) | 20.35 |
| $R_{\text{free}}$ (%)** | 24.96 |
| Monomers / a.u. | 2 |
| No. of residues | 325 |
| No. of atoms | 2721 |
| Protein | 2502 |
| Water | 219 |
| R.m.s. deviation |  |
| Bond lengths (Å) | 0.018 |
| Bond angles (°) | 2.040 |
| Mean B value (Å<sup>2</sup>) |  |
| Protein | 31.28 |
| Water | 35.59 |

\*Values in parentheses are for the highest resolution shell.
\*\*Throughout the refinement, 5% of the total reflections were held aside for $R_{\text{free}}$ .

**Table S2.**
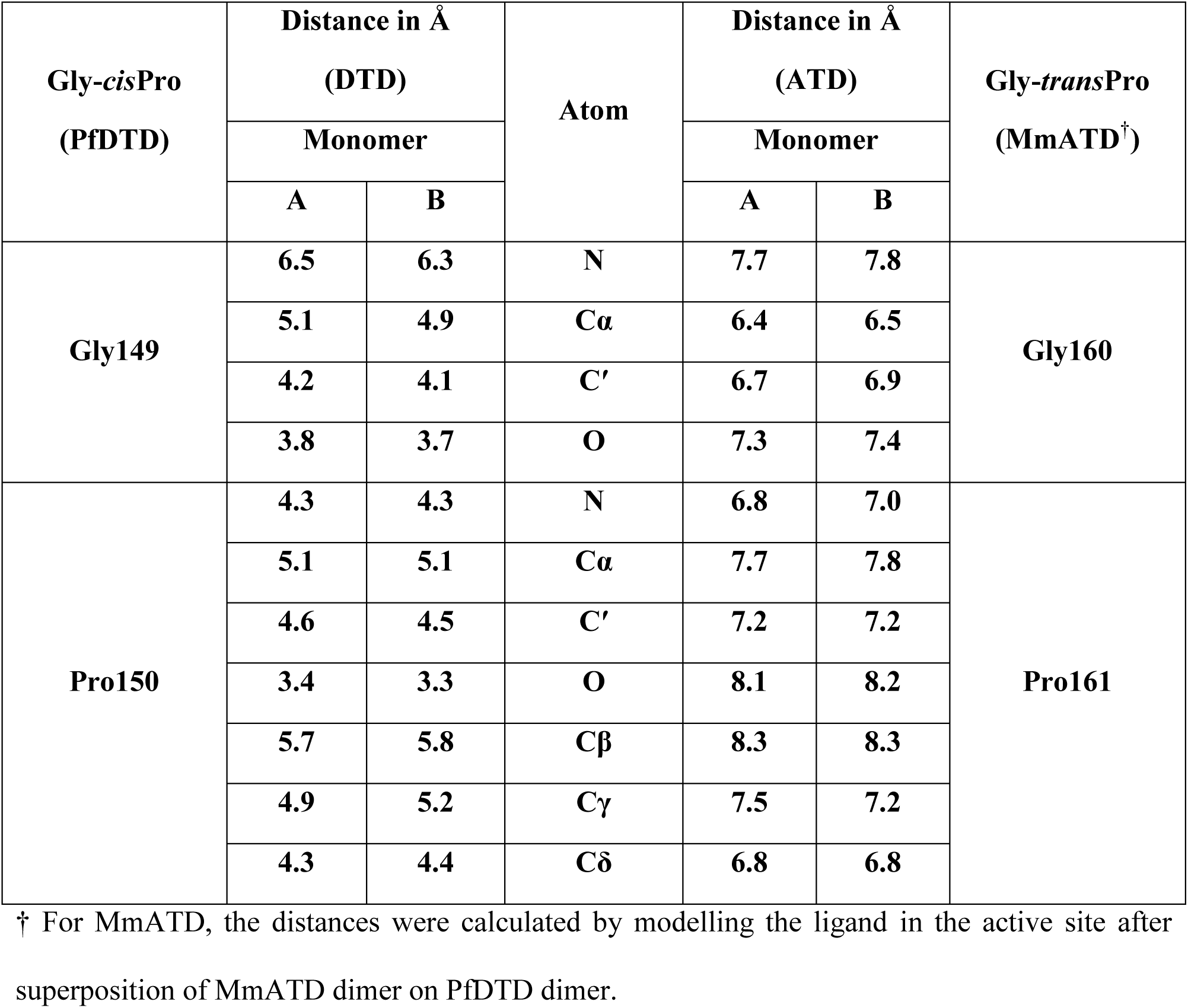
“Additional” space in ATD’s active site pocket. Comparison of distances between atoms of Gly-Pro residues and Cα of the ligand D-Tyr3AA for PfDTD (PDB id: 4NBI) and MmATD†.

**Table S3.** Comparative analysis of enrichment of tRNA^Thr^(G4•U69) genes and tRNA^Cys^(G4•U69) genes in representative organisms. Relative abundance of tRNA^Thr^(G4•U69) genes and tRNA^Cys^(G4•U69) genes in representative organisms.

| Organism | tRNA<sup>Thr</sup> genes |  |  | tRNA<sup>Cys</sup> genes |  |  |
| --- | --- | --- | --- | --- | --- | --- |
|  | Total | G4•U69 | % | Total | G4•U69 | % |
| *Strongylocentrotus purpuratus* * | 57 | 14 | **24.6** | 31 | 0 | **0** |
| *Latimeria chalumnae* | 22 | 5 | **22.7** | 21 | 2 | **10** |
| *Danio rerio* | 722 | 209 | **28.9** | 144 | 1 | **0.7** |
| *Takifugu rubripes* | 28 | 12 | **42.9** | 12 | 0 | **0** |
| *Xenopus tropicalis* | 245 | 74 | **30.2** | 83 | 2 | **2.4** |
| *Gallus gallus* | 10 | 4 | **40.0** | 10 | 0 | **0** |
| *Rattus norvegicus* | 16 | 4 | **25.0** | 40 | 0 | **0** |
| *Mus musculus* | 17 | 4 | **23.5** | 57 | 1 | **1.8** |
| *Pan troglodytes* | 18 | 4 | **22.2** | 27 | 0 | **0** |
| *Homo sapiens* | 20 | 4 | **20.0** | 29 | 1 | **3.4** |

\* *Strongylocentrotus purpuratus* is the only organism known so far that does not belong to phylum Chordata (it belongs to phylum Echinodermata) but shows enrichment of tRNA<sup>Thr</sup>(G4•U69) genes.

**Movie 1. The flip from Gly-*cis*Pro in DTD to Gly-*trans*Pro in ATD.** Movie depicting the remodeling of the local network of interactions due to *cis*-to-*trans* switch.

**Data 1. List of organisms whose genomes have been sequenced, highlighting the presence or absence of ATD.**

